## Supplementary information for "Stochastic model of T Cell repolarization during target elimination (II)"

### Stochastic model of T Cell repolarization during target elimination (II) SUPPLEMENTARY MATERIAL

#### 1 Dynein attachment during the first few seconds

When $\beta < \pi$ only a fraction of MTs intersect the center of the IS, visually demonstrated in Fig. S1, and their number is given by the ratio $q_{\text{CIS}}$, see Fig. 3c in the main text. Considering the number of MT intersecting with the center of the IS, the number of attached dyneins can be estimated using Eq. 3 in the main text.

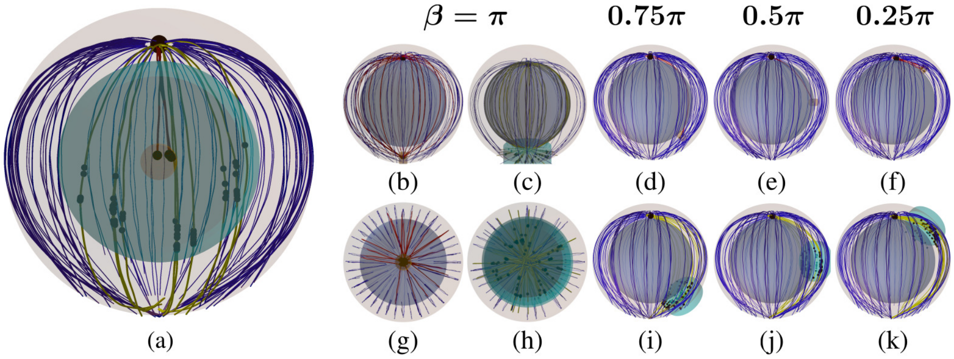


Fig. S1: Snapshots from the beginning of simulations with different angles $\beta$ between the IS and the initial position of the MTOC, sketched in Fig. 1a in the main text. The plasma membrane and the nucleus are represented by the transparent outer and inner sphere, respectively. The cyan and small brown cylinder denote the complete IS and its center, respectively. Unattached(blue), capture-shrinkage(red) and cortical sliding(yellow) MTs sprout from the MTOC depicted by the big black sphere. Small black spheres represent attached dyneins. (a) Snapshot from the beginning of the simulation with $\beta = 0.5\pi$ from the perspective of the IS. Just a fraction of MTs pass through the IS and its center. (b)(g) Cell with the center of the IS from the horizontal and vertical perspective, respectively. (c)(h) Cell with the whole IS from the horizontal and vertical perspective, respectively. (d)(e)(f) Cells with the center of the IS. (i)(j)(k) Cells with the whole IS.

It can be seen in Figs. S2(a-f) that the number of attached dyneins initially sharply increases due to the fast attachment of the dyneins. As the time progresses, dyneins detach and the number of attached dyneins decreases and subsequently approaches the estimated values. The number of dyneins is lower than the estimated value when $\rho_{\text{IS}} = 500\mu\text{m}^{-2}$, which can be explained by the fact that the stronger force results in a faster movement and bigger opposing forces increase the probability of the dynein detachment. Moreover, the dynein force presses the MTOC stronger against the nucleus, see Figs. S4(d-f).

The estimates of attached dyneins are lower when $\beta > 0.8\pi$, see Figs. S2(g-i), since only a fraction of MTs are long enough to reach the IS. As was stated in the first part of our work (1), the number of MT beads is uniformly distributed between 15 and 20. Consequently, only $66, 33, 17\%$ of MTs are long enough to reach the IS when $\beta = 0.8, 0.9, 0.95\pi$, respectively. The number of attached dyneins are lower than estimates when $\beta > 0.5\pi$, see Figs. S2(g-i), due to the fact that the nucleus intersects the path of the MTOC to the IS, as visualized in Figs. S1b-d, and the repulsive force from the nucleus increases the dynein detachment probability.

The number of attached dyneins differ from the estimates for the cases $\beta = 0.15\pi$ and $\beta = \pi$, see Figs. S2j and k. When $\beta < 0.25\pi$ the cytoskeleton is immediately dragged to the IS due to the short initial MTOC-IS distance and dyneins detach, see Figs. S4c and f. In the case of $\beta = \pi$, every MT with the sufficient length intersects the center of the IS, visualized in Figs. S1b and g. Around 20% of MTs have a sufficient length to reach the IS in the first seconds of the simulation. The number of dyneins estimated by the Eq. 3 in the main text would be greater than the number of dyneins in the IS. To avoid this contradiction, we assume that all dyneins attach during the first seconds of the simulation. It can be seen in Fig. S2k that the measured number of attached dyneins is substantially lower. It can be explained by the fact that dyneins act on MTs sprouting in all orientation from the IS, visualized in Figs. S1b and g, and therefore act in competition and mutually increase force-dependent detachment rate (1).

#### 2 Additional results for one IS: $\beta$-dependence for different mechanisms

##### 2.1 Capture-Shrinkage mechanism

The supplementary Movie S8 shows the MTOC repositioning under the effect of the capture-shrinkage mechanism, $\rho_{\text{IS}} = 300\mu\text{m}^{-2}$, $\beta = 0.75\pi$. The process is also visualized for the case of $\beta = 0.75\pi$ in Fig. S3. At the beginning of the simulation, only a small fraction of MTs intersects the IS and they sprout from the MTOC in one direction (visualized

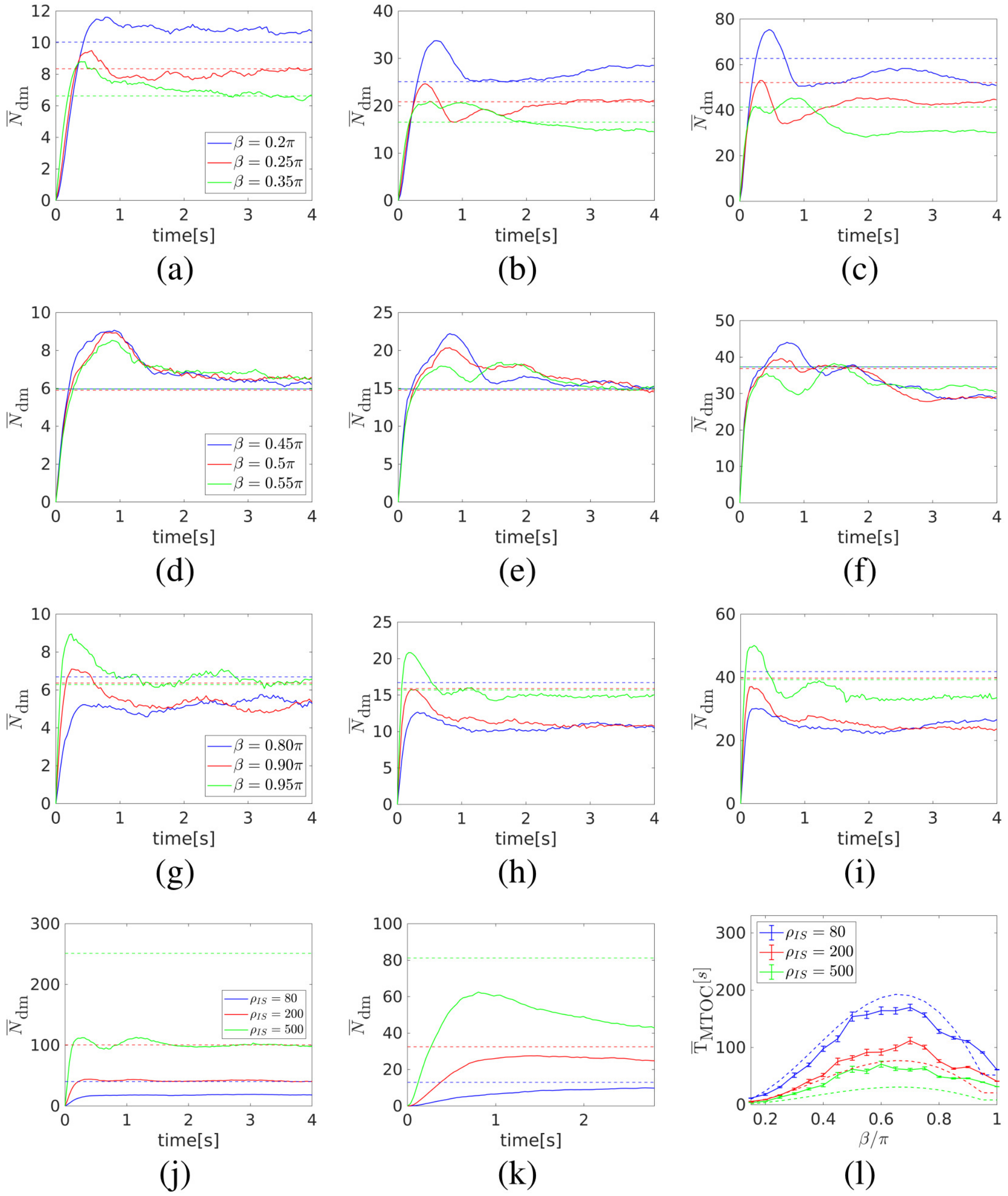


Fig. S2: The dependence of the average number of dyneins $\overline{N}_{\rm dm}$ on the time during the first seconds of repositioning. The dashed lines represent estimations. (a, d and g) $\rho_{\rm IS} = 80\mu{\rm m}^{-2}$. (b, e and h) $\rho_{\rm IS} = 200\mu{\rm m}^{-2}$. (c, f and i) $\rho_{\rm IS} = 500\mu{\rm m}^{-2}$. (j) $\beta = \pi$ (k) $\beta = 0.15\pi$ (l) Comparison of the measured(solid) and estimated(dashed) times of the repositioning.

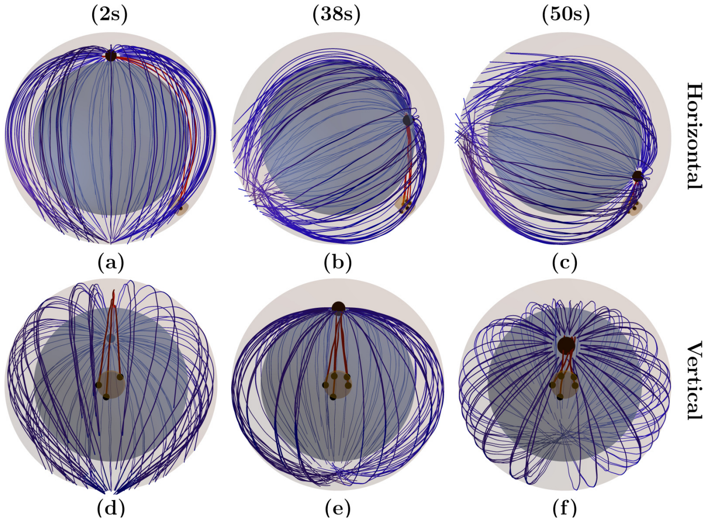


Fig. S3: Snapshots from the time evolution of the spindle configuration under the effect of the capture-shrinkage mechanism (dynein density $\rho_{\rm IS} = 300\mu{\rm m}^{-2}$, initial angle between MTOC and IS $\beta = 0.75\pi$). MTs connect to the MTOC represented by the large black sphere. Blue and red curves represent unattached and attached MTs, respectively. The brown cylinder depicts the center of the IS, where the capture-shrinkage dyneins represented by small black spheres are located. (a and d) $d_{\rm MIS} \sim 8.0\mu$m. A stalk connecting the MTOC and IS is formed. (b and e) $d_{\rm MIS} \sim 4\mu$m. The stalk shortens and the MTOC approaches the IS. (c and f)$d_{\rm MIS} \sim 1\mu$m. The MTs in stalk depolimerized and MTOC is in the close proximity of the IS.

Fig. S4: Capture-shrinkage mechanism: (a-c) Dependence of the average MTOC-IS distance $\bar{d}_{\rm MIS}$ on time. Dependencies of the average MTOC velocity $\bar{v}_{\rm MTOC}$ in insets (a-c), the average number of dyneins acting on microtubules $\bar{N}_{\rm dm}$ (d-f) and the MTOC-center distance $\bar{d}_{\rm MC}$ in the insets of (d-f) on the average MTOC-IS distance. Black dashed lines denote transitions between different phases of the repositioning process. (g)(h)(i) Probability densities of the angle $\alpha$ between the first MT segments and the direction of the MTOC movement for the dynein density $\rho_{\rm IS} = 300\mu{\rm m}^{-2}$ at the beginning of the simulation. (g and h) Time $t$ = 10s. (i) $t$ = 5s.

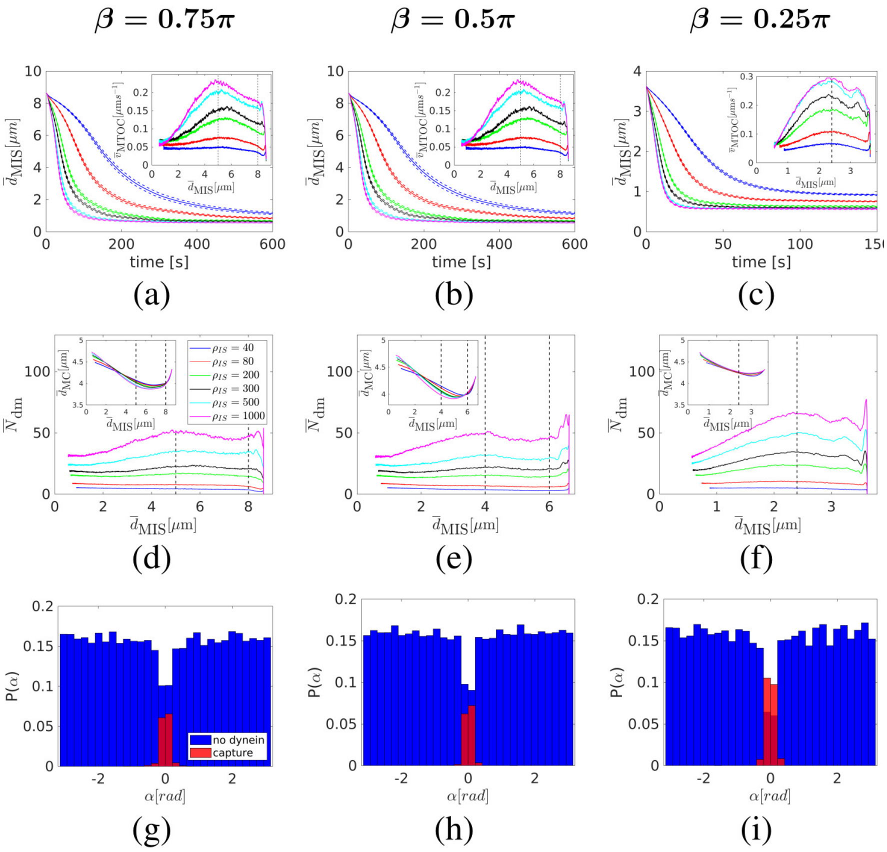


in Figs. S3a and d. The MTs attach to dynein at the beginning of the simulation and form a narrow stalk connecting the MTOC and the IS. As the MTOC approaches the IS, unattached MTs are pushed back by the friction forces and the spindle "opens", visually demonstrated in Figs. 3e and f. As the time progresses, the stalk of MTs shortens and the MTOC approaches the IS, see Figs S3b, c, e and f.

When $\beta > 0.25\pi$, the process can be divided into three phases based on the time evolution of the MTOC velocity, see Figs. S4a and 4b. In the first phase comprising approximately the first $0.5\mu$m of the MTOCs trajectory, the velocity increases and then in some cases decreases to a local minimum. In the second phase, the velocity slightly increases until it reaches a maximum and then in the third phase, it declines again. By comparison of Figs. S4ab and S4de one realizes that the time evolution of the velocity is determined by the time evolution of the number of attached dyneins, which can be understood by considering the forces from the nucleus. At the beginning of the simulations, MTs intersects the IS (visually demonstrated in Figs. S3a and d) leading to the dynein attachment and the formation of a narrow MT stalk, see Figs. S4g-i, visually Fig. S3a and d. As the MTOC moves to the IS in the direction given by the MT stalk, it is pressed against the nucleus, see Figs. S4d and e, visually demonstrated by Figs. S1 and S3. The opposing forces of the nucleus repel the MTs resulting in the detachment of dyneins, see Figs. S4d and e, because the detachment rate is force dependent. The force from the nucleus decreases as the MTOC slides on its surface, causing the slight increase of the MTOC velocity despite the constant pulling force (compare Figs. S4ab and S4de). The number of attached dyneins remains approximately constant in the second phase, see Figs. S4d and e, since unattached MTs in opening cytoskeleton are pushed back by friction forces and unlikely to attach to dynein, visualized in Figs. S3d-f. The decrease of the number of attached dynein in the third phase can be explained by the shortening of the MTs in the stalk due to the depolymerization lowering the probability of dynein attachment. More importantly, the detachment probability increases due to the opposing force of the cytoskeleton being dragged from the nucleus to the membrane, see Figs. S4d and e. Consequently, the repositionings have similar characteristics as the one in the cell when $\beta = \pi$ (1).

When $\beta = 0.25\pi$ the repositioning has only two phases as can be seen in Fig. S4f, since the velocity of the MTOC rises quickly and then continuously decreases. Due to the short initial MTOC-IS distance the nucleus does not present an obstacle and MTOC is pulled directly to the cell membrane, see Fig. S4f. Therefore, the number of attached dyneins decreases due to the same causes as in the third phase when $\beta > 0.25\pi$, compare Figs. S4de and f.

#### 2.2 Cortical Sliding mechanism

The repositioning process under the effect of the cortical sliding mechanism is visually demonstrated in Fig. S5 and in the supplementary Movie S9, $\tilde{\rho}_{\rm IS} = 300\mu{\rm m}^{-2}$, $\beta = 0.75\pi$. In contrast to the capture-shrinkage, MTs attach to the cortical sliding dynein in the whole IS, which is substantially larger than its center. Figs. S5a and d shows that the MTOC is pulled to the IS by MTs aligned in a large stalk. As the MTOC approaches the IS, dyneins detach and remaining attached dyneins are increasingly located at the periphery of the IS, visually demonstrated in Figs. S5b and e. Almost all dyneins detach at the end of the repositioning, see Figs. S5c and f.

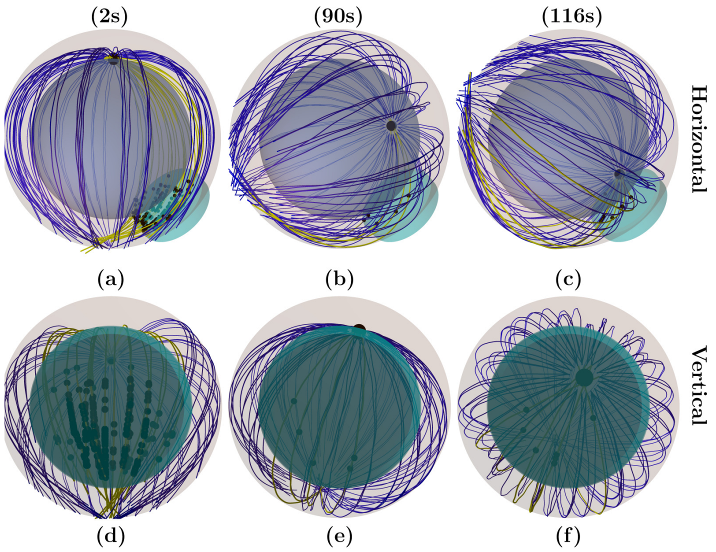


Fig. S5: Snapshots from the time evolution of the MT cytoskeleton under the effect of the cortical sliding mechanism (dynein density $\rho_{\text{IS}} = 300\text{m}^{-2}$, $\beta = 0.75\pi$). The cyan cylinder denotes the area of the IS. Blue and yellow lines represent unattached and attached MTs, respectively. The small black spheres in the IS stand for the positions of dyneins attached to MTs. (a and d) $d_{\text{MIS}} \sim 8.0\mu\text{m}$. A stalk connecting the MTOC and IS is formed. The attached MTs are sprouting from the MTOC in one direction since the beginning. (b and e) $d_{\text{MIS}} \sim 7.5\mu\text{m}$. The number of attached dyneins decreased and motors are attached in one half of the IS and mainly at its periphery. (c and f) $d_{\text{MIS}} \sim 1\mu\text{m}$. The MTOC is in the close proximity of the IS. Dyneins are attached to MTs solely on the periphery of the IS.

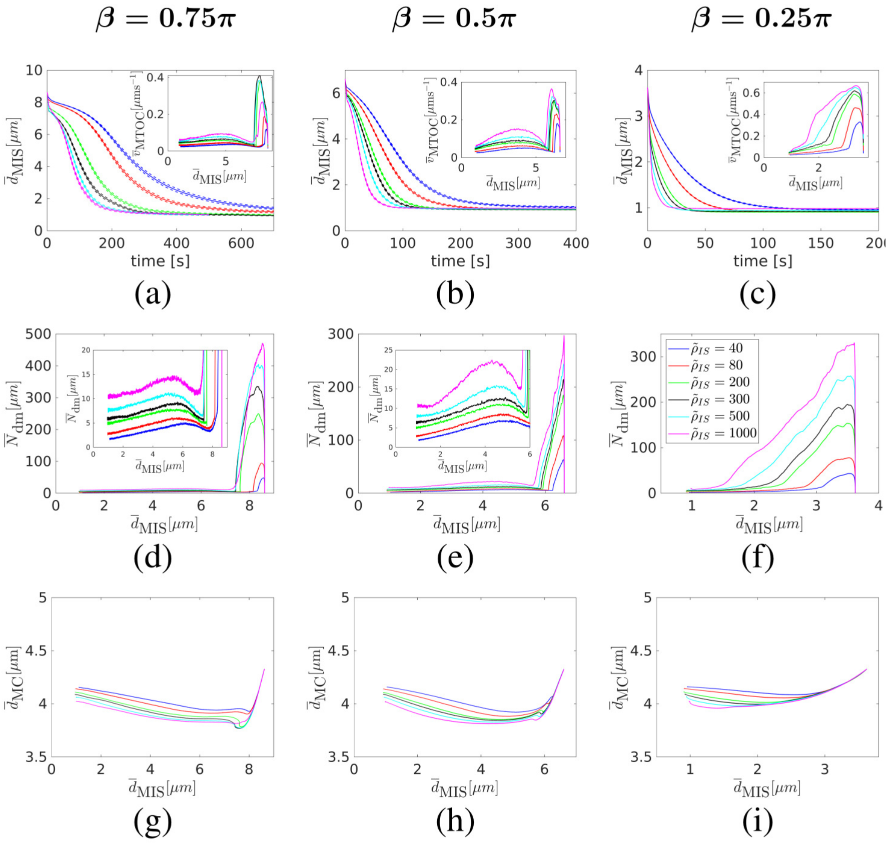


Fig. S6: Cortical sliding mechanism. (a-c) The dependence of the average MTOC-IS distance $\bar{d}_{\text{MIS}}$ on the time is shown. Inset: the dependence of the average MTOC velocity $\bar{v}_{\text{MTOC}}$ on the average MTOC-IS distance are shown. The dependencies of the average number of dyneins acting on MTs $\bar{N}_{\text{dm}}$(d-f), the average distance between the MTOC and the center of the cell $\bar{d}_{\text{MC}}$(g-i) on the average MTOC-IS distance are shown.

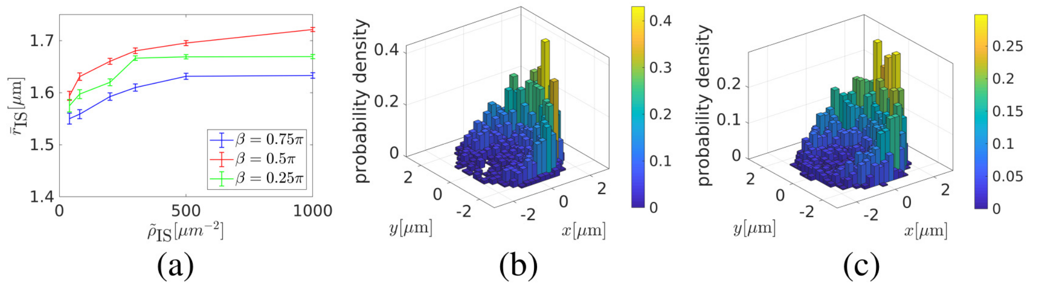


Fig. S7: Attached cortical sliding dyneins. (a) The dependence of the average distance $\bar{r}_{\mathrm{IS}}$ of attached cortical sliding dyneins from the axis of the IS on dynein area density $\tilde{\rho}_{\mathrm{IS}}$ is shown. (b-c) The two-dimensional probability density of attached dyneins in the IS in the middle of the repositioning, $\overline{d}_{\mathrm{MIS}} = 3\mu\mathrm{m}$, $\beta = 0.5\pi$, is shown. (b) $\tilde{\rho}_{\mathrm{IS}} = 80\mu\mathrm{m}^{-2}$. (c) $\tilde{\rho}_{\mathrm{IS}} = 1000\mu\mathrm{m}^{-2}$.

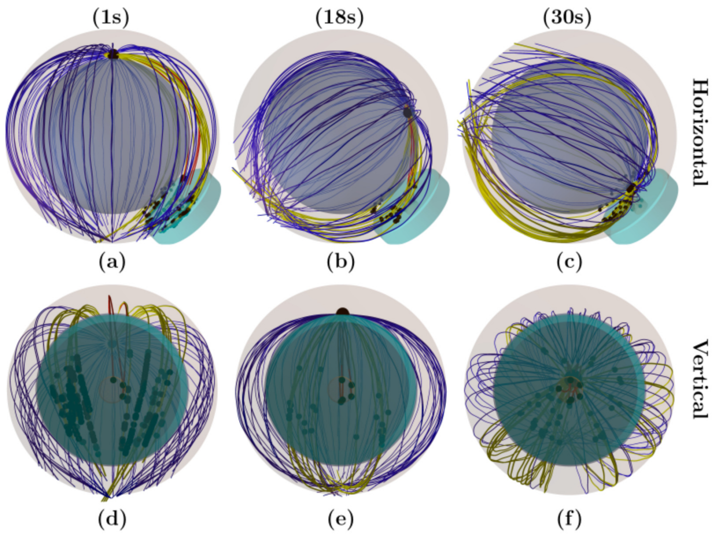


Fig. S8: Snapshots from the time evolution of the MT cytoskeleton under the combined effect of both mechanisms (dynein density $\rho_{\mathrm{IS}} = \tilde{\rho}_{\mathrm{IS}} = 300\mu\mathrm{m}^{-2}$, $\beta = 0.5\pi$). The cyan and brown cylinders denote the IS and its center, respectively. Blue, yellow and red lines represent unattached, cortical sliding and capture-shrinkage MTs, respectively. The small black spheres in the IS stand for the positions of the dyneins attached to MTs. (a and d) $d_{\mathrm{MIS}} \sim 8\mu\mathrm{m}$. A stalk of capture-shrinkage MTs connecting the MTOC and the center of the IS is formed at the beginning. Cortical sliding MTs attach at the periphery of the IS and do not intersect the center of the IS. (b and e) $d_{\mathrm{MIS}} \sim 4\mu\mathrm{m}$. The capture-shrinkage MTs in the stalk shorten and the MTOC approaches the IS. The MTOC slides on the surface of the nucleus. (c and f) $d_{\mathrm{MIS}} \sim 1\mu\mathrm{m}$. The MTs in the stalk are depolymerized and the MTOC is in the close proximity of the IS. The MTOC detached from the nucleus and moves to the membrane. Cortical sliding dyneins are attached at the IS periphery.

Similarly to the previous case of the capture-shrinkage mechanism, the repositioning can be divided into three phases when $\beta > 0.25\pi$. Contrarily, the transition points between the phases depend on the density $\tilde{\rho}_{\mathrm{IS}}$, see Figs. S6a and b. The time evolution of the velocity of the MTOC is determined by the number of dyneins, compare Figs. S6a and b with Figs. S6d and e. The stalk of MTs connecting the MTOC and the IS is formed at the beginning of the simulation, visually in Figs. S5a and d, and the dynein quickly attach to MTs. Due to the opposing force of the nucleus the number of dynein decreases during the first phase, see insets of Figs. S6d and e. The decrease is steeper than in the previous case, compare Figs. S4d and e with S6d and e, due to the fact that the cortical sliding lacks a firm anchor point characteristic for the capture-shrinkage. In the second phase, the number of dynein rises, see insets of Fig. S6d and e, since the opposing force of the nucleus decreases as the MTOC slides on the surface. The number of attached dyneins decreases in the third phase because the MTOC does not recede substantially from the nucleus of the cell, see Figs. S6g-i, indicating that the MTs do not follow the membrane resulting in the lower attachment probability of dynein, visually demonstrated in Figs. S5c and f.

The MTOC displays only one phase when $\beta = 0.25\pi$, see Figs. S6c and f because of short initial distance and the fact that the MTOC is not an obstacle on the MTOC's journey, visualized in Figs. S1f and k. The number of attached dynein decreases from the beginning of the simulation, see Fig. S6f, since the MTs do not copy the cell membrane as the MTOC approaches the IS.

Fig. S7 shows that similarly to the case of $\beta = \pi$ (1) the attached dyneins are located predominantly at the periphery of the IS and their average distance from the axis of the IS slightly increases with the density. Consequently, we report that attached dyneins are predominantly located on the periphery of the IS regardless of the initial configuration of the cell.

#### 2.3 Combined mechanisms

The supplementary Movie S10 shows the MTOC repositioning under the combined effects of both mechanism, $\rho_{\text{IS}} = \tilde{\rho}_{\text{IS}} = 300\mu\text{m}^{-2}$, $\beta = 0.5\pi$. At the beginning of the simulation, MTs intersecting the center of the IS attach to capture-shrinkage dyneins and form a narrow stalk connecting the MTOC and the IS, visually demonstrated in Figs. S8a and d. Cortical sliding dynein attach and pull MTs in the periphery of the IS. Consequently, the MTs in the stalk shorten and the MTOC slides on the surface of the cell towards the IS, visually Figs. S8b and e. At the end of the repositioning, the MTOC recedes from the nucleus and is pulled to the center of the IS by capture-shrinkage MTs, visually Figs. S8c and f.

We examine the combined effect of the mechanisms in two initial configurations: $\beta = 0.5, 0.75\pi$. When $\beta = 0.25\pi$ the repositioning is already very fast, see Figs. S4 and S6 and Fig. 4b in the main text, and the combination is not needed for an efficient repositioning. The interplay of the two mechanisms when $\beta = \pi$ was previously analyzed in the first part of our work (1). Figs. S9a and b shows that the combination of the two mechanisms always leads to a faster repositioning. Moreover, two mechanisms with relatively small dynein densities can outperform the dominant mechanism with a substantially larger density. For example when $\beta = 0.75\pi$, the capture-shrinkage mechanism with the density $\rho_{\text{IS}} = 1000\mu\text{m}^{-2}$ is slower than any combination of mechanisms where $\tilde{\rho}_{\text{IS}}, \rho_{\text{IS}} \geq 200\mu\text{m}^{-2}$, see Fig. S9a. It generally holds for both angles $\beta$ that the combination of two mechanisms when $\tilde{\rho}_{\text{IS}}, \rho_{\text{IS}} \geq 200\mu\text{m}^{-2}$ is faster than an individual mechanism.

The capture-shrinkage mechanism was identified as the dominant mechanism when $\beta = 0.75\pi$. Fig. S9a shows that the repositioning gets faster with the rising cortical sliding density. Moreover, the average number of attached capture-shrinkage dyneins increases with the cortical sliding density, see Fig. S9d. This can be explained by the alignment of cortical sliding MTs in the direction of the MTOC movement, see Fig. S9f. In the case when capture-shrinkage acts alone, the unattached MTs are unlikely to attach to capture-shrinkage, since they are pushed back by friction forces, visually demonstrated in Fig. S3. MTs attached to cortical sliding tend to align to the MTOC movement, see Figs. S9f and i and visually demonstrated by the Fig. S8. Due to the alignment they are not pushed back by friction forces and they are more likely to be captured in the narrow center of the IS. Figs. S9f and i shows that the dominant central peak of the cortical sliding MTs is interrupted in the middle by the narrow peak of capture-shrinkage MTs proving that cortical sliding MTs are passed to the capture-shrinkage mechanism. Therefore, the cortical sliding mechanism gives capture-shrinkage substantial advantage compared to the case when capture-shrinkage acts alone by increasing the probability that the MT will attach to dyneins in the center of the IS.

Furthermore, the cortical sliding helps capture-shrinkage dyneins because they share the load from opposing forces. The sharing of forces leads to the decrease of the force dependent detachment probability and to the increase of attached capture-shrinkage dyneins. To summarize, cortical sliding supports capture-shrinkage by increasing the attachment probability by aligning the MTs with the MTOC movement and by decreasing the detachment probability by sharing the load from opposing forces.

Fig. S9e demonstrates that when the density of cortical sliding dyneins is fixed, the repositioning gets faster with the increasing capture-shrinkage density. Moreover, the number of attached cortical sliding dyneins increases with the rising capture-shrinkage density mainly at the end of the repositioning. This may come as a surprise, since the presence of capture-shrinkage decreases the number of cortical sliding MTs due to their attachment in the center of the IS, see Fig. S9f and i. Moreover, dyneins in the part of the IS behind the center(from the MTOC's perspective) cannot attach to MTs, see Fig. S9c, since they are distant from any filament. Fig. 9c clearly demonstrates that the attached cortical sliding dyneins form interesting ”slashed” patterns indicating that the dyneins behind IS remain unattached. Seemingly, the capture-shrinkage mechanism harms the cortical sliding by stealing the MTs and by preventing it to reach a part of the IS.

However, the capture-shrinkage dynein pull in alignment towards the center of the IS and provide a firm anchor point that the cortical sliding is otherwise missing. Figs. S6g-i clearly demonstrates that the MTOC does not substantially recedes from the nucleus in the case of the sole cortical sliding. Consequently, MTs do not copy the cell membrane and the dynein attachment probability decreases. The pulling force of capture-shrinkage dyneins causes that the membrane is followed more closely in the case of combined mechanisms, see Figs. S9g-h, resulting in the increase of the number of attached cortical sliding dyneins, see Fig. S9e. Additionally, capture-shrinkage dyneins share the load from opposing forces leading to the decrease of the detachment probability of cortical sliding dyneins and therefore to the bigger number of attached dyneins.

To conclude, the cortical sliding mechanism supports capture shrinkage by passing the MTs. Capture-shrinkage promotes cortical sliding by providing it a firm anchor point. The two mechanisms support each other by sharing the load from opposing forces. To conclude, the mechanisms act in a fascinating synergy in every configuration of the cell.

Fig. S9: Combination of capture-shrinkage and cortical sliding mechanisms: (a and b) Dependence of the averaged times $\overline{\mathrm{T}}_{\mathrm{MTOC}}$ (MTOC-IS distance $d_{\mathrm{MIS}} < 1.5\mu$m) on the capture-shrinkage dyneins density $\rho_{\mathrm{IS}}$. Cortical sliding densities are given by the color. (c) The two-dimensional probability density of attached dyneins in the IS is shown, $\beta = 0.75\pi$, $\tilde{\rho}_{\mathrm{IS}} = \rho_{\mathrm{IS}} = 500\mu\mathrm{m}^{-2}$, the MTOC-IS distance $\mathrm{d}_{\mathrm{MIS}} \sim 3.5\mu$m. (d) Dependence of the average MTOC-IS distance $\overline{\mathrm{d}}_{\mathrm{MIS}}$ on time. In the inset: Dependence of the average number of attached capture-shrinkage dyneins $\overline{N}_{\mathrm{capt}}$ on the average MTOC-IS distance $\overline{d}_{\mathrm{MIS}}$. $\beta = 0.75\pi$, $\rho_{\mathrm{IS}} = 500\mu\mathrm{m}^{-2}$. The cortical sliding density $\tilde{\rho}_{\mathrm{IS}}$ is given by the color. (e) Dependence of the average MTOC-IS distance on time. In the inset: Dependence of the average number of attached cortical-sliding dyneins $\overline{N}_{\mathrm{cort}}$ on the average MTOC-IS distance. $\beta = 0.5\pi$, $\tilde{\rho}_{\mathrm{IS}} = 500\mu\mathrm{m}^{-2}$. The capture-shrinkage density $\rho_{\mathrm{IS}}$ is given by the color. (f) The probability density of the angle $\alpha$ between the first MT segment and the direction of the MTOC motion is shown. $\tilde{\rho}_{\mathrm{IS}} = \rho_{\mathrm{IS}} = 200\mu\mathrm{m}^{-2}$, $t = 10$s, $\overline{\mathrm{d}}_{\mathrm{MIS}} \sim 4\mu$m, $\beta = 0.5\pi$. (g and h) The dependence of the average MTOC-center distance $\overline{d}_{\mathrm{MC}}$ on the average MTOC-IS distance $\overline{d}_{\mathrm{MIS}}$. $\tilde{\rho}_{\mathrm{IS}} = 40\mu\mathrm{m}^{-2}$. Capture-shrinkage density $\rho_{\mathrm{IS}}$ is given by the color. (i) The probability density of the angle $\alpha$ between the first MT segment and the direction of the MTOC motion is shown. $\tilde{\rho}_{\mathrm{IS}} = \rho_{\mathrm{IS}} = 200\mu\mathrm{m}^{-2}$, $t = 10$s, $\overline{\mathrm{d}}_{\mathrm{MIS}} \sim 8\mu$m, $\beta = 0.75\pi$.

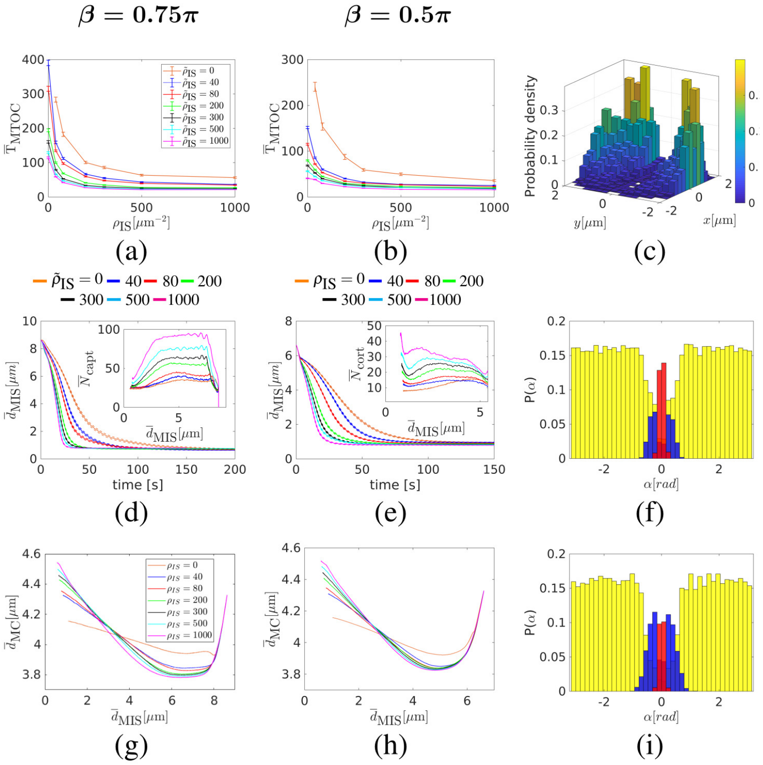
